# Impact of a Cancer-Associated Mutation on Poly(ADP-ribose) Polymerase1 Inhibition

**DOI:** 10.1101/2024.11.13.623412

**Authors:** Neel Shanmugam, Shubham Chatterjee, G. Andrés Cisneros

## Abstract

Poly(ADP-ribose) polymerase1 (PARP1) plays a vital role in DNA repair and its inhibition in cancer cells may cause cell apoptosis. In this study, we investigated the effects of a PARP1 variant, V762A, which is strongly associated with several cancers in humans, on the inhibition of PARP1 by three FDA approved inhibitors: niraparib, rucaparib and talazoparib. Our work suggests that these inhibitors bind to the V762A mutant more effectively than to the wild-type (WT), with similar binding free energies between them. Talazoparib inhibition uniquely lowers the average residue fluctuations in the mutant than the WT including lower fluctuations of mutant’s N- and C-terminal residues, conserved H-Y-E traid residues and donor loop (D-loop) residues which important for catalysis more effectively than other inhibitions. However, talazoparib also enhances destabilizing interactions between the mutation site in the HD domain in the mutant than WT. Further, talazoparib inhibition significantly disrupts the functional fluctuations of terminal regions in the mutant, which are otherwise present in the WT. Lastly, the mutation and inhibition do not significantly affect PARP1’s essential dynamics.

**GRAPHICAL ABSTRACT:** Figure.
Impact of V762A mutation on inhibition of PARP1.

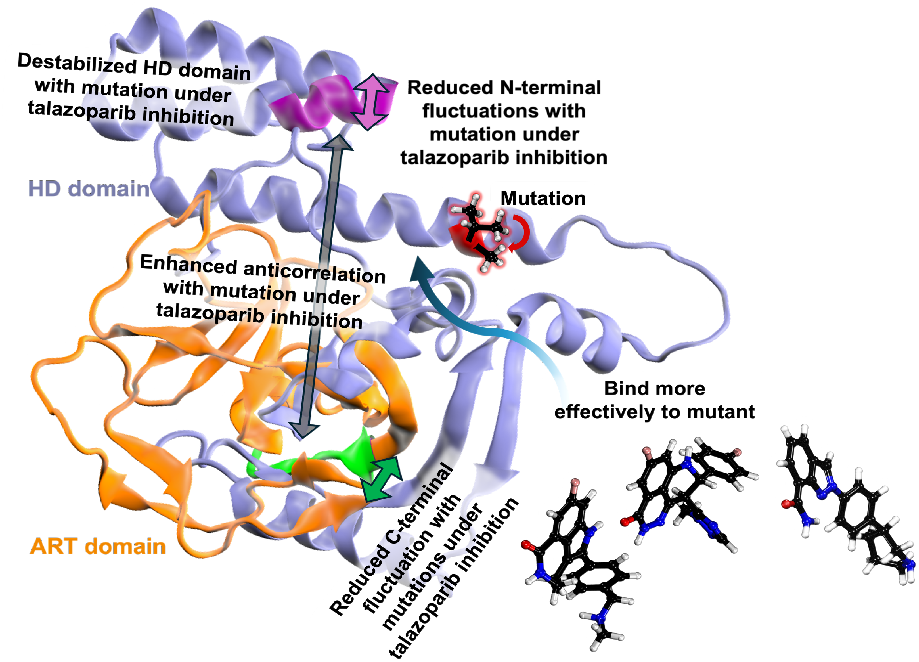

## INTRODUCTION

DNA damage occurs at a rate of approximately 1 million changes per cell each day in humans,(1) caused by factors like such as UV radiation, ionizing radiation, chemical agents, errors during replication, and cellular metabolic processes.(2–4) DNA damage can contribute to diseases, with the accumulation of cancer-causing mutations in DNA playing a key role in tumor development and cancer progression.(5, 6) Humans have developed several DNA damage response (DDR) pathways including base excision repair (BER), non-homologous end joining (NHEJ), homologous recombination (HR), and Okazaki-fragment processing pathways to minimize this genomic instability.(7)

Poly(ADP-ribose) polymerase1 (PARP1) is a DNA binding protein, member of the PARP family, which consists of at least 18 distinct proteins in humans.(8) PARP-1 is responsible for 85%-90% of poly(ADP-ribosylation) (PAR) activity.(9–15) PARP1 plays a part in all above-mentioned DNA repair pathways, and is a key enzyme in the BER pathway. For BER, PARP1 recruits repair proteins to single-stranded breaks (SSBs), facilitating DNA repair. While PARP1 does not directly stimulate HR, cells deficient in HR are highly sensitive to PARP1 inhibition. During DNA replication, PARP1 detects SSBs that naturally occur when Okazaki fragments on the lagging strand are processed, recruiting repair proteins such as XRCC1. Additionally, PARP1 influences NHEJ, a repair mechanism for double-strand breaks (DSBs). (7, 16–23)

Upon sensing DNA damage, PARP-1 undergoes conformational changes and initiates PARylation to add Poly(ADP-ribose) (PAR) chains covalently to itself or any other target proteins like histones, DNA polymerases, DNA ligases, etc as shown in **Figure 1(B) to 1(E)**. This modification recruits DNA repair proteins, such as XRCC1 and Lig-IIIα, to the damaged sites to facilitate DNA repair.(24–29) PARP1 is a multi-domain enzyme as shown in **Figure 1(A)** it is structured into six regions, each with distinct functions. The N-terminal region of PARP1, functioning as the DNA-binding domain (DBD), consists of three zinc finger motifs: ZN-1 (residues 1-100), ZN-2 (residues 101-215), and ZN-3 (residues 216-370). These motifs collaborate to detect DNA strand breaks, with ZN-2 playing a critical role in this interaction.(30, 31) The central region which is the auto modification domain (AD) includes a BRCT domain (371 - 500). The BRCT domain acts as the acceptor site for auto-poly(ADP-ribosyl)ation (auto-PARylation). The C-terminal region houses the catalytic domains namely, tryptophan-glycine-arginine (WGR, residues 501-650) domain whose function is not fully understood but is implicated in DNA interactions, helical subdomain (HD, residues 651-785) and the ART (residues 786-1014) subdomain.(32–34). The ART subdomain contains several highly conserved residues — H862, Y896 and E988 and the donor loop (D-loop, residues P881 to Y889) of the active site of PARP1. H862 and Y896 are crucial for NAD^+^ binding to PARP1 and its for polymerase’s activity. The D-loop helps shape the binding site and interacts with NAD^+^. (35)

**Figure 1.**
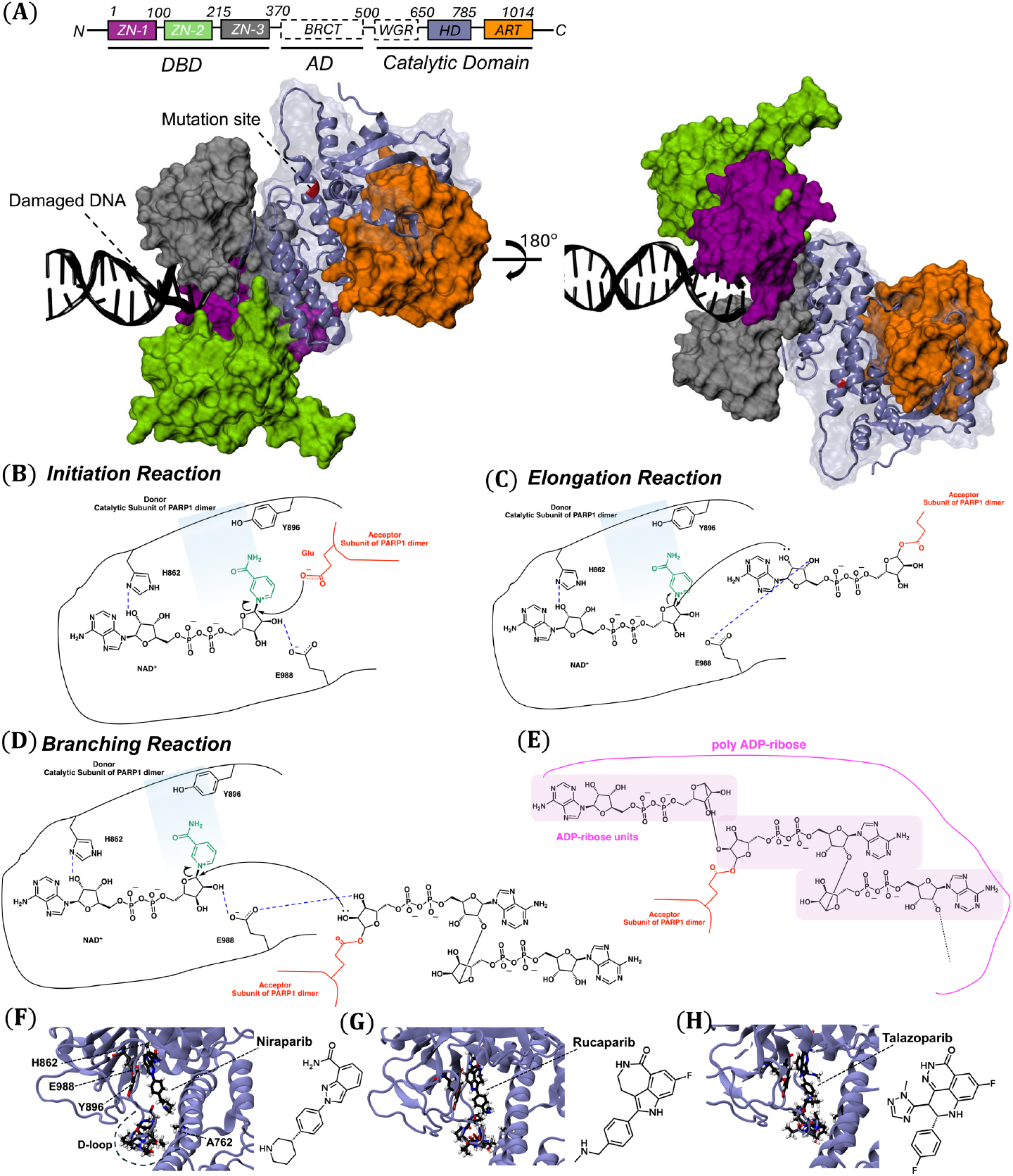
**(A)** Structural and multi-domain organization of near full-length PARP-1 in complex with a DNA double-strand break (PDB ID: 4DQY(104)). **(B-D)** Mechanism of poly-ADP-ribosylation by PARP-1(99–101) **(E)** poly ADP-ribose consisting of ADP-ribose units. **(F-H)** show the active sites of mutated PARP1 PDBs (4R6E, 6VKK, and 4UND respectively) with niraparib, rucaparib and talazoparib inhibitors. The active site consists of H-Y-E traid and D-loop residues. 2-D structures on the side panels depict the structures of the respective inhibitors.

PARP1 inhibitors compete with NAD^+^ for binding in the active site (**Figure 1(F) to 1(H)**).(36) In the absence of an inhibitor, PARP is recruited to a SSB site for DNA. With an inhibitor in the NAD^+^ binding site, PARP1 may lose the ability to activate PARP-dependent repair pathways and cannot dissociate from DNA due to inhibited catalytic activity and/or direct trapping, resulting in replication fork (RF) stalling during DNA replication and, ultimately, stalled RFs collapsing to form double-strand breaks (DSBs).(37, 38)

In a BReast CAncer gene 1/2 (BRCA1/2) proficient cells, DSBs can be repaired via HRR. By contrast, PARP inhibition for BRCA-deficient cancer cells can lead to cell cycle arrest and apoptosis. This approach, which is also known as synthetic lethality, is exploited therapeutically by using PARP inhibitors for eliminating BRCA-mutant cancer cells. (32, 35, 39–52)

After the N-terminal region of PARP1’s ZN-1 domain binds to damaged DNA sites, there is an initiation of allosteric signaling event that activates the catalytic function at catalytic domain in C-terminal. This process is crucial for PARP’s role in DNA repair.(53) Some inhibitors, in addition to blocking catalysis, also trap PARP1 and PARP2 on damaged DNA, forming cytotoxic complexes that are more effective at killing cancer cells. This “PARP trapping,” or “PARP poisoning,” varies among inhibitors and is independent of their catalytic inhibition potency, highlighting the dual mechanism of action of these drugs.(54–59)

Murai and colleagues demonstrated that bulky PARP inhibitors, such as olaparib, can trap the PARP-1/DNA complex. Their findings support the hypothesis that allosteric reverse signaling occurs, where the CAT domain sends signals back to the N-terminal DNA binding domain, disrupting its normal function. This trapping mechanism has significant implications for the therapeutic targeting of PARP in cancer treatments.(60–63)

In a study by Zandarashvili et al., PARP1 inhibitors were classified into three categories based on their allosteric effects on PARP1: type I (EB-47 and BAD), type II (talazoparib and olaparib), and type III (rucaparib, niraparib, and veliparib).(36) In addition to catalytic inhibition, type I inhibitors exert a strong allosteric effect, causing instability in the HD domain causing allosteric changes that enhance interactions with the DNA break and slow down its release. By contrast, type II inhibitors have minimal impact on PARP-1 allostery, resulting in only slight increases in affinity for the DNA break. Type III inhibitors stabilize the HD domain and promote release from the DNA break.

PARP1 has been reported to show alterations in multiple cancers.(64) Moreover, a single PARP1 variant has been shown to affect response to PARP1 inhibition.(65) We have previously reported a common single-nucleotide polymorphism (SNP) in the PARP-1 gene, rs1136410,(66) that results in the V762A PARP1 variant. This mutation is strongly associated with lung cancer, ovarian cancer, and other neoplastic diseases in humans. Experimental validation confirmed that the mutation leads to overactivation of the enzyme, impairing its ability to repair DNA breaks.(66)

In the present work, we performed molecular dynamics (MD) simulations to investigate the possible impact of V762A on the inhibition of PARP1 by three FDA-approved inhibitors: niraparib(67–70), rucaparib(71–73) and talazoparib(74, 75), sold under the brand names Zejula, Rubraca, and Talzenna, respectively. The organization of the paper is as follows: the next section describes the details of MD simulations of the inhibited systems, and MD simulation calculations for binding free energy calcualtions. Subsequently, we present the details for post simulation analysis: root mean square deviation (RMSD), root mean square fluctuation (RMSF), principle components of motion, and movement correlation matrices, and energy decomposition analaysis (EDA), followed by concluding remarks.

## MATERIAL AND METHODS

### PARP1 Simulations

The crystal structures for the catalytic domain of the WT PARP1 dimer, both without an inhibitor (PDB ID: 4ZZZ(76)) and bound to talazoparib (PDB ID: 4PJT(77)) were were obtained from the RCSB Protein Data Bank. For the mutated dimer, structures without an inhibitor (PDB ID: 5WS1) and bound to niraparib (PDB ID: 4R6E(77)), rucaparib (PDB ID: 6VKK(78)), and talazoparib (PDB ID: 4UND(79)) were also obtained. Each dimer was converted into a monomer by isolating chain A. UCSF Chimera(80) was then used to align the mutated PARP-1 structures bound to niraparib and rucaparib with the WT structure without an inhibitor to generate PDB files for WT structure bound to niraparib and rucaparib.

MD simulations were carried out via AMBER18’s(81) pmemd.cuda(82). Inhibitor parameters were generated using the Antechamber tool in AmberTools20. (82) The inhibitors were initially minimized independently (without PARP-1) using Sander. Eight systems were prepared: WT PARP-1 without an inhibitor, with niraparib, with rucaparib, and with talazoparib, as well as mutated (V762A) PARP-1 without an inhibitor, with niraparib, with rucaparib, and with talazoparib. Each system was neutralized using chloride counterions and solvated in a TIP3P(83) water box extending a minimum distance of 10 Å between the edge of the protein to the edge of the box using the LEaP(81) module of AmberTools20 and the ff14SB(84). The systems were first minimized with 50 steps of steepest descent and 950 steps of conjugate gradient, applying positional restraints of 500 kcal mol−^1^ to hold the enzyme in place. This was followed by a second minimization with positional restraints reduced to 10 kcal mol−^1^. The systems were then heated to 300K using Langevin dynamics(85–87) with a collision frequency of 2 ps^−1^ and then equilibrated for 15 nanoseconds under NPT conditions, during which positional restraints were gradually removed. Finally, production simulations were conducted on these unrestrained systems for 500 nanoseconds each, resulting in a total of 1.5 microseconds of simulation time across the three replicas with an integration time-step of 2 fs, and trajectories were saved every 2 ps. Long-range Coulombic interactions were handled under periodic boundary conditions with the smooth particle mesh Ewald (sPME)(88) method using a 10 Å cutoff for nonbonded interactions.

Relative binding free energies between the WT and mutant structures under inhibition were calculated with independent trajectories using thermodynamic integration (TI).(89) **Figure 2** shows the thermodynamic cycle that was considered when calculating relative binding free energies. The free energy was calculated using the equations below:

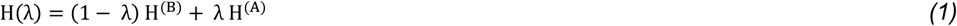

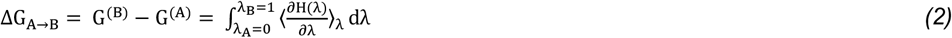

**Figure 2.**
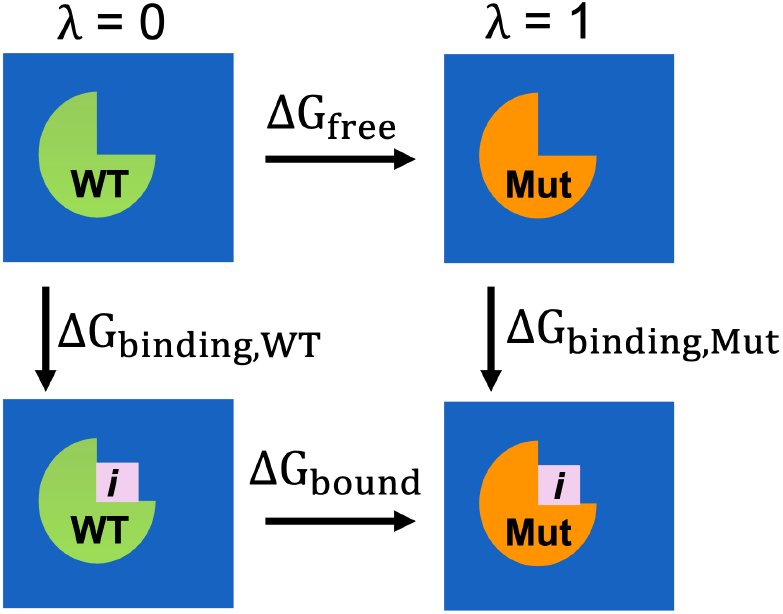
Thermodynamic cycle for a relative binding free energy calculation of the respective inhibitor (*i*) in wild type (WT) and mutant (Mut) systems. The vertical arrows show inhibitor binding to the protein (WT and mutant). The horizontal arrows show the alchemical transformations between WT and Mut (V762 -> A762).

Here, ΔG is the free energy difference for a segment, H(λ) is the Hamiltonian of the system, dependent upon λ which is a parameter that varies from the initial state A where λ = 0 to the final state B where λ = 1 as shown in equation (1), and where denotes ⟨. . . ⟩_λ_ an ensemble average at a particular λ value, which is obtained in practice from MD simulations. Equation 2 was integrated numerically using the trapezoidal rule with simulations of 30 ns run at discrete values of λ using default soft-core non-bonded potentials(90) in Amber19. In this study for calculating the relative binding free energy (ΔΔG_binding,WT → Mut_)f the inhibitors between WT and mutant, we used WT and mutant as the initial and final states respectively as indicated in **Figure 2** and **equation (3)**.

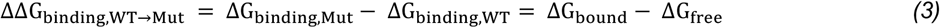

The TI calculations for λ values were carried under the same procedure as for the MD described above.

### Post-simulation analysis

The backbone root mean squared deviation (RMSD), residue-wise root mean squared fluctuation (RMSF), principal component, and dynamic cross-correlation analyses (PCA and DCC respectively) were calculated using AMBER’s CPPTRAJ(91) program. The .nmd files that visualize the principal components are included in the Zenodo repository. Hydrogen bonds (H with N, O, and F) involving the inhibitor that lasted for a substantial portion of production (> 10%) were analyzed using CPPTRAJ by calculating the distance and energy between atoms over time. Normal modes were visualized using the Normal Mode Wizard in VMD.(92–99) The first 8 PCA modes were calculated from each trajectory using the final 10,000 snapshots from the production. (93)

Energy decomposition analaysis (EDA) was performed for the entire production ensemble to investigate the non-bonded (NB) inter-molecular interactions between two fragments using Amber_EDA available on GitHub.(100–102) This method calculates the average inter-molecular interaction energy, 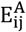 between every pair of fragments. The differences in these interactions between various systems are then determined using equation (4).

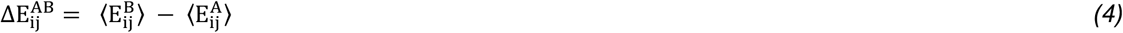

Here, system A represents the reference system, and system B is the target system. The term 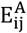 denotes the average non-bonded interaction energy between fragment i in system A with all other fragments j in system A, with a similar interpretation for system B. The difference in non-bonded intermolecular interactions between systems A and B, 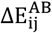 (henceforth 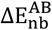) is calculated by averaging over three replicates of each system. In this context, the WT system was used as the reference for all the calculations.

## RESULTS

We calculated the time dependent RMSD for all α carbon (Cα) atoms in the catalytic domain of PARP1 to investigate the difference in deviations of PARP1 backbone and residues in WT and mutant systems (**Figure SI1**). The RMSD remained below 3 Å for most of the simulation time for both WT and mutant systems, indicating equilibrated protein structures during the production segment of the simulations. Further, the average deviation of the HD domain in the WT structures was higher under talazoparib inhibition compared to niraparib and rucaparib inhibitions (**Figure 3(A)**). As shown in **Figure 3(B)**, the average change in deviation between WT and mutant PARP1 was higher in the mutated structures for the apo system. By contrast, under niraparib and rucaparib inhibition, the average deviation of the Cα in the PARP1 backbone was minimally affected. For the talazoparib-inhibited system, these deviations decreased with the V762A mutation.

**Figure 3.**
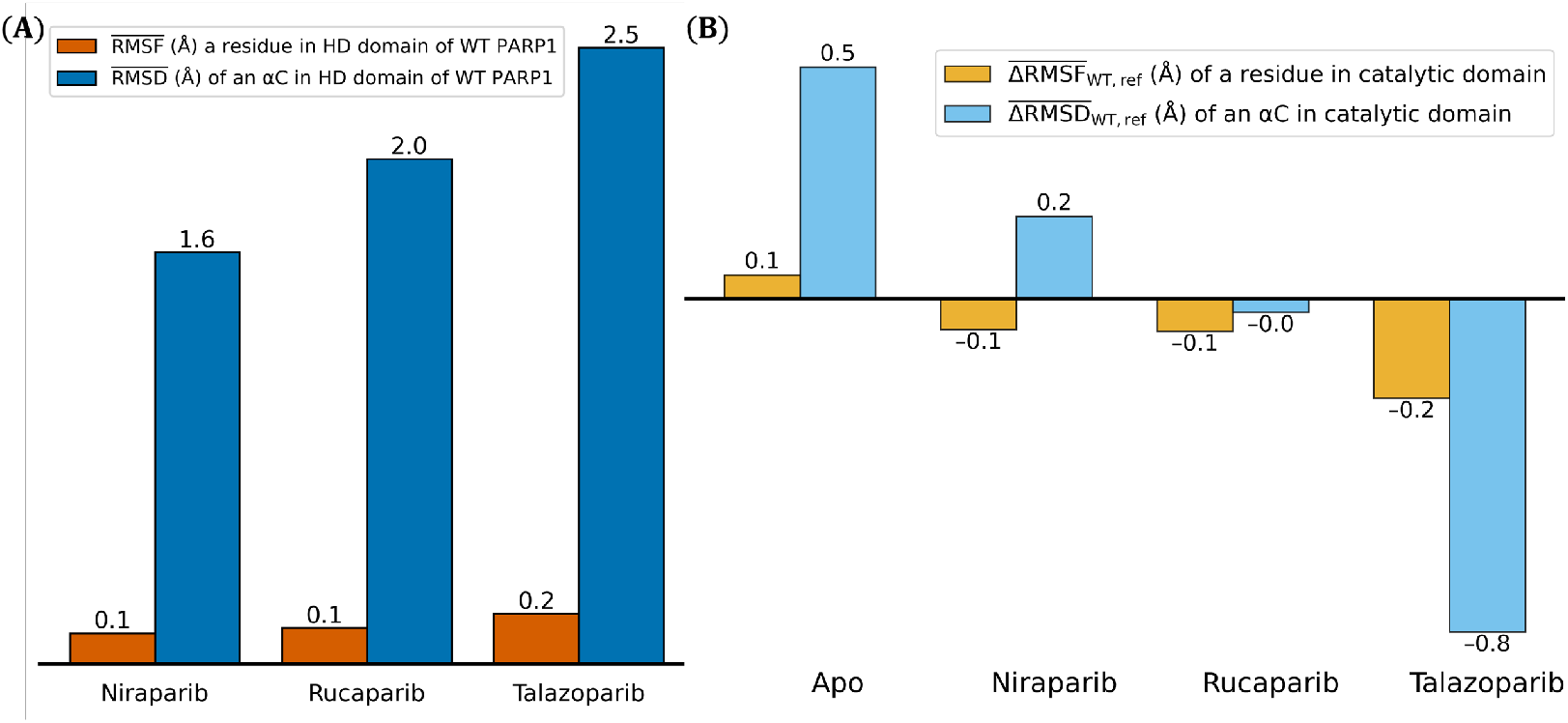
**(A)** Plots of average fluctuations in Å of a residue in HD domain of WT PARP1 (in orange) and plots of average deviation in Å of an Cα in PARP1 backbone in HD domain (in blue) **(B)** Plots of difference in average change in fluctuations in Å of a residue between WT and mutant systems (in orange) and plot of difference in the average deviation in Å of an Cα in PARP1 backbone between WT and mutant systems (in skyblue). In both cases, WT PARP1 serves as the reference.

The average change in fluctuations for individual residues in the HD domain for the WT structures was slightly higher under talazoparib inhibition compared with niraparib and rucaparib inhibition (**Figure 3(A)**). Higher RMSD and RMSF of Cα atoms and residues respectively in the HD domain of WT PARP1 during talazoparib inhibition, compared to niraparib and rucaparib inhibitions, suggest that niraparib and rucaparib provide greater stabilization of the HD domain in WT structures than talazoparib. This observation is consistent with the experimental results reported by Zandarashvili.(36) Further, the average residual fluctuations were minimally affected under no inhibition, niraparib inhibition and rucaparib inhibition. However, for the talazoparib-inhibited system, these deviations decreased with the mutation (**Figure 3(B)**).

**Figure 4(A)** to **(D)** shows a heat map of the difference in average residual fluctuations. In the case of apo the residual fluctuations for the majority of residues increased for the mutant. By contrast, the majority of residues showed decreased fluctuations in inhibited mutated systems than the respective inhibited WTs. In all inhibitor-bound systems, the C-terminal residues of the mutant remained more stable compared to the WT, with this stability being particularly pronounced in the talazoparib-inhibited system (**Figure 4(A)** to **(D)**). The N-terminal residues in the mutant also showed significantyly higher stability under talazoparib inhibition compared to other inhibitions. Further, when inhibited by rucaparib, the N-terminal residues exhibited greater fluctuations in the mutant than the WT. As shown in **Figure 4(E)**, the mutation caused negligible changes in the average fluctuation of H862. However, the average fluctuations of Y896 and E988 increased with the mutation in the apo system, remain almost unaffected in the presence of niraparib or rucaparib, and decreased with the mutation in the presence of talazoparib. Unlike what is observed for the H-Y-E triad, the average fluctuations of the D-loop-residues changed significantly with the mutation under inhibition. The fluctuations decreased upon inhibition, with the greatest reduction observed for talazoparib, followed by niraparib, and the least reduction for rucaparib (**Figure 4(F)**).

**Figure 4.**
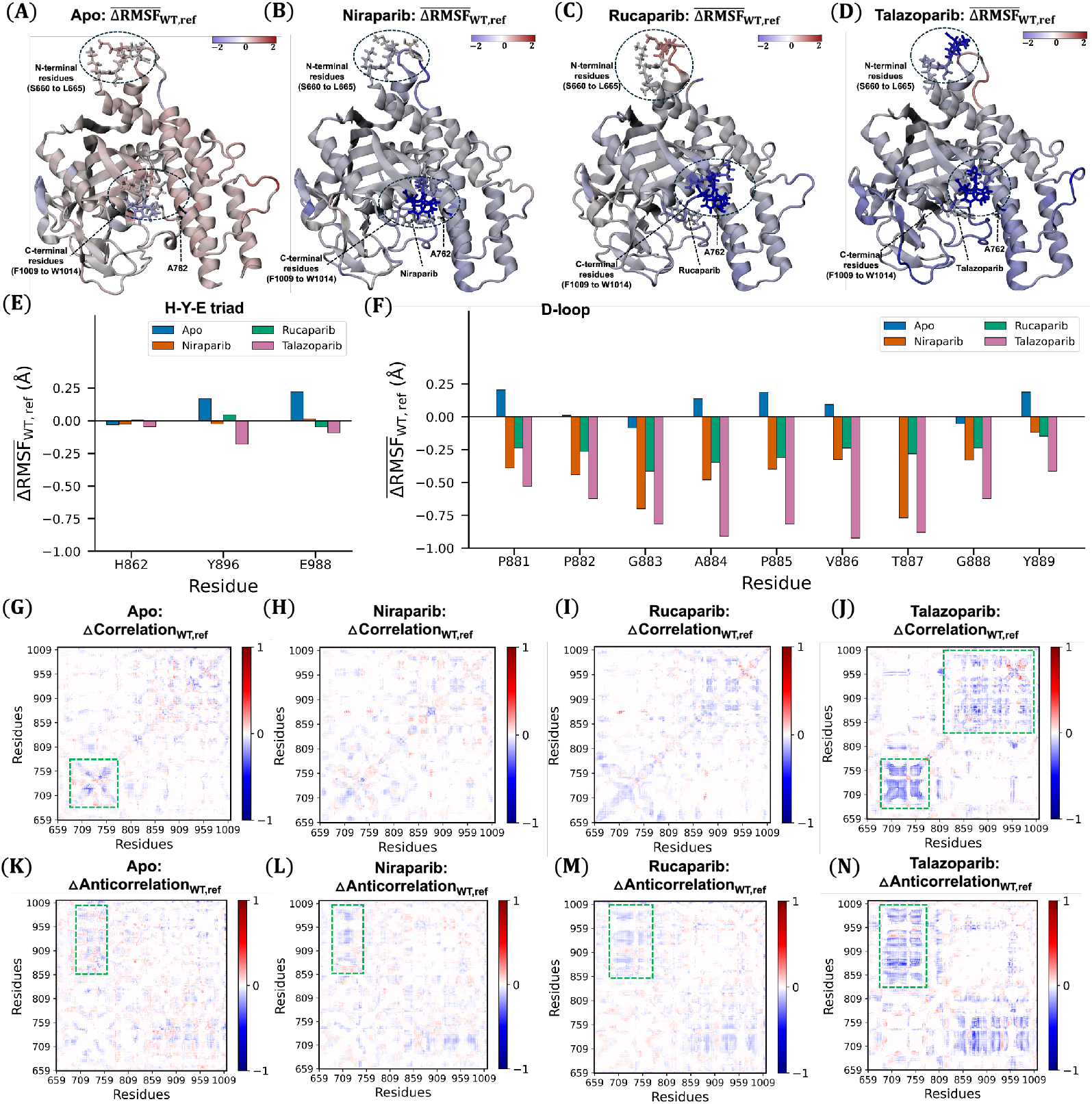
**(A-D)** Average difference in RMSF in Å between WT and mutant systems, a red(blue) shift indicates increased(decreased) fluctuations. **(E**,**F)** Average difference in RMSF in Å of for the H-Y-E traid and D-loop residues respectively. **(G-J)** Dynamic cross-correlation (DCC) difference plots of correlated motion between WT and mutant in apo systems, niraparib-inhibited, rucaparib-inhibited and talazoparib-inhibited systems respectively. **(K-N)** DCC difference plots of anticorrelated motions of residues between WT and mutant in apo systems, niraparib-inhibited, rucaparib-inhibited and talazoparib-inhibited systems respectively.

Dynamic cross-correlation (DCC) analysis was performed to investigate the effect of the mutation on the relative motion of the subdomains of PARP1 (**Figure 4(G)** to **(N)**). In the case of apo (**Figure 4(G)**) higher correlated motion is observed among the N-terminal residues in the WT compared to the mutant. **Figure 4(K)** suggests that the motion of N- and C-terminal residues are slightly more anticorrelated in the WT-than the mutant-apo system. Similarly, in inhibited systems, the N- and C-terminal residues exhibit higher anti-correlated motion in the WT system compared to the mutant. However, under inhibition, the anticorrelation is much higher than the apo systems with talazoparib-inhibition showing the highest level of anti-correlated movement between the terminals, as shown in **Figures 4(L),(M) and (N)**. Further, under talazoparib inhibition, the movement of residues in the N- and C-terminal regions is more strongly correlated within their respective regions (N-terminal with N-terminal, C-terminal with C-terminal) in the WT compared to the mutant as shown in **Figure 4(J)**.

The essential dynamics of the domains are illustrated by the normal mode analysis (NMA). The top 8 modes among 100 modes calculated were considered (**Figure SI6**). The predominant motion in every case is a combination of breating and rocking motion in all the WT and mutant systems of apo and inhibition (movie in Zenodo repository).

The inhibitor can influence the interactions between the protein residues and the mutation site, potentially stabilizing or destabilizing the mutated region. To investigate this possibility we conducted an energy decomposition analysis (EDA). We observe that all the systems apart from rucaparib-inhibition show reduced inter-molecular interactions with the mutation site, with the talazoparib inhibited system showing the highest destabilizing interactions with the mutation site. As shown in **Figure 5**, residues in the binding pocket show significant changes in interactions with the mutation site. In the case of apo, Q759 and Y710, located in the binding pocket significantly destabilize and stabilize the mutation site respectively. In the case of niraparib-inhibition, Y710 again shows significant stabilizing interactions with the mutation site. Residues D766 and the D-loop residue P885 significantly destabilize the mutation site. In the case of inhibition with rucaparib, the change in residual interactions with the V762A mutation is minimal. However, in the case of talazoparib-inhibition, we observe that the mutation site is significantly destabilized by the surrounding residues in the binding site. A D-loop residue Y889 majorly destabilize the mutation site.

**Figure 5.**
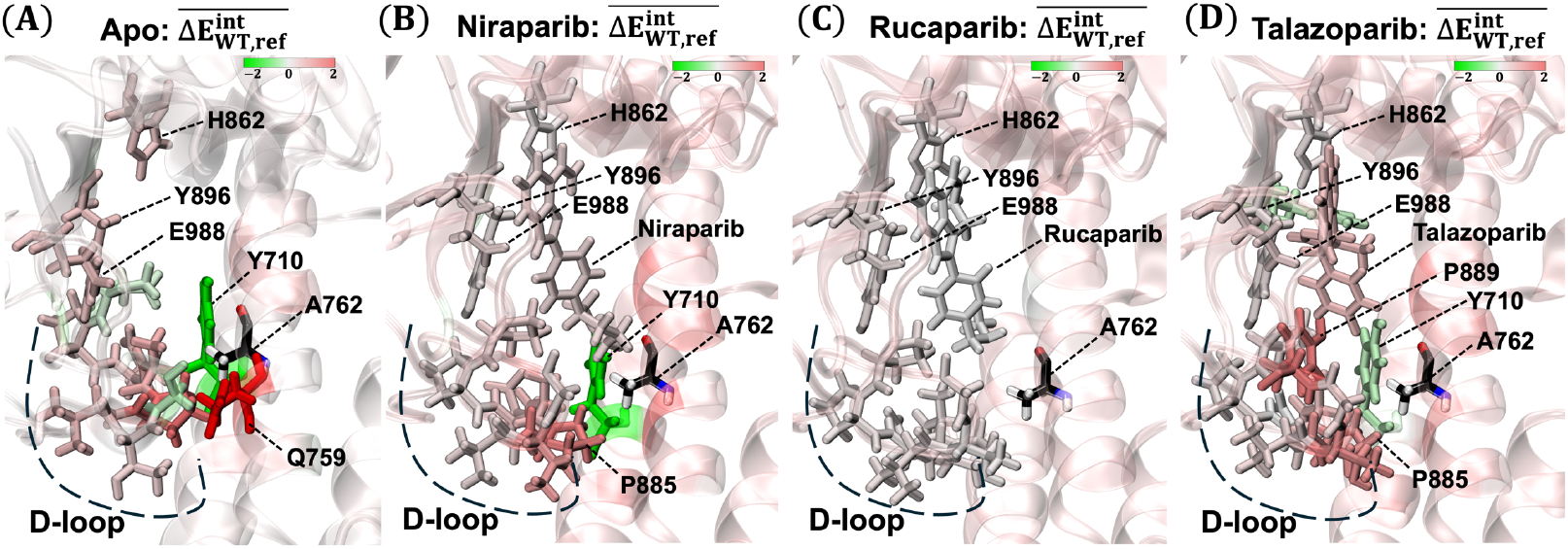
Heat maps showing the differences in non-bonded interaction energies between PARP1 residues and the respective inhibitor with the mutation site (V762 in WT and A762 in the mutant) for the WT and mutant under different conditions: **(A)** without inhibition, **(B)** with niraparib inhibition, **(C)** with rucaparib inhibition, and **(D)** with talazoparib inhibition. In all cases, the WT serves as the reference. For simplicity, only important residues in the binding pocket are shown. Color scale ranges from green to red, representing higher stabilizing to destabilizing interaction energies with A762 than V762.

As shown in **Figure 6**, the overall change in interaction energies between the HD domain residues and the mutation site was lower under niraparib and rucaparib inhibition than under talazoparib. Additionally, as noted earlier, the HD domain in the WT structures showed a greater average deviation under talazoparib inhibition compared to niraparib and rucaparib (**Figure 3(A)**). As reported by Zandarashvili et al.(36), stabilization of the HD domain in PARP1 corresponds to faster release from a DNA break. This suggests that, under niraparib and rucaparib inhibition there could be a faster release of PARP1 from DNA than the WT, compared to that under talazoparib inhibition.

**Figure 6.**
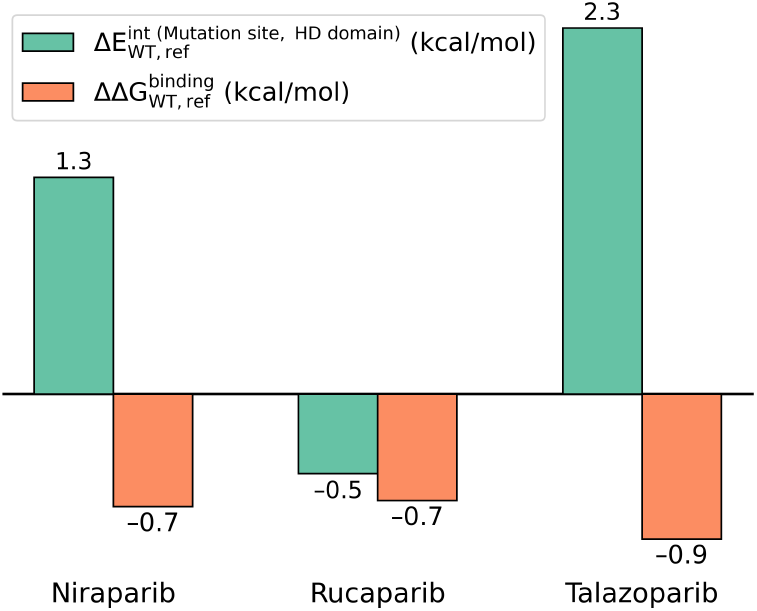
Change in average interaction energy (green) between the HD domain and the mutation site (A762 in the WT and V762 in the mutant) under niraparib, rucaparib, and talazoparib inhibition, using WT as the reference. Relative average binding free energies (orange) of niraparib, rucaparib, and talazoparib to PARP1, comparing the WT to the mutant, with the WT as reference.

Next, to understand how the mutation affects the binding to the inhibitors to the active site we performed TI calculations. PARP1 backbone remained stable during the simulations (**Figure SI4(A)** to **(F)**) suggesting equilibrated PARP1 structure during calculations. Considerable overlap of phase space at neighboring λ values (**Figure SI4(G)** to **(L))** indicate statistically correlated structures for neighboring λ values. 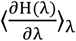 values for each λ are presented in **Figure SI5**. Negative relative binding free energies (**Figure 6**) for the inhibitors indicate that all three inhibitors show slightly improved binding to the PARP1 cancer variant than the WT.

## DISCUSSION and CONCLUSION

The mutant system showed a significantly lower deviation in the backbone from the crystal structure under talazoparib inhibition compared with the other inhibitors (**Figure 3(B)**). This suggests a more stable catalytic domain for the mutant than the WT under talazoparib inhibition. C-terminal residues of the mutant fluctuated less than those of the WT in both apo and inhibited systems (**Figure 4(A) to 4(D)**), with lowest fluctuations under talazoparib inhibition. This suggests that talazoparib most strongly affects the functional motion of the C-terminal when the mutation is present. These results are consistent with experimental reports from Zandarashvili et al., (36) which have indicated that allosteric reverse signaling may occur in PARP1, where the catalytic domain sends signals to the N-terminal DNA-binding domain, disrupting its typical function. Similar to C-terminal residues, we observed least fluctuating N-terminal residues in the mutant than the WT under talazoparib inhibition than other inhibitors (**Figure 4(D)**). To our knowledge, the role of N-terminal residues in the catalytic domain of PARP1 remains unexplored. Current observations, therefore, present new opportunities for further investigation of these residues.

Of the three inhibitors studied, talazoparib makes the HD domain significantly more unstable in the mutant than in the WT (**Figure 6**). According to Zandarashvili et al.(103), stabilization of the HD domain in PARP1 is linked to faster release from DNA breaks. Therefore, our findings suggest that under talazoparib inhibition, the mutant may retain damaged DNA more effectively than the WT, compared to other inhibitors. Among the conserved H-Y-E residues, H862 fluctuations were similar in both WT and mutant across all systems, while Y896 and E988 showed consistent fluctuations between WT and mutant in the apo, niraparib-, and rucaparib-inhibited systems but decreased more significantly under talazoparib inhibition (**Figure 4(E)**). Additionally, the greater stability of D-loop residues (**Figure 4(F)**) in the mutants of the inhibited systems, especially with talazoparib, suggests that these inhibitors stabilize the D-loop more effectively in mutants than in WTs.

Under inhibitor conditions, the level of anti-correlated movement between the N- and C-terminal regions is significantly higher than in the apo state, with talazoparib showing the strongest anti-correlation, as seen in **Figures 4(K),(L), (M), and (N)**. Additionally, talazoparib inhibition results in stronger correlations within each terminal region (N-terminal with N-terminal, C-terminal with C-terminal) in the WT compared to the mutant, as shown **in Figure 4(J)**. This indicates that talazoparib inhibition substantially disrupts the functional fluctuations of these terminal regions in the mutant, which are preserved in the WT. The primary motion of PARP1 in both apo and inhibited systems is a combination of breathing and rocking motion. Our free energy simulation results suggest that all three inhibitors considered have stronger binding to mutant than WT, with similar relative free energy relative free energy affinities.

Overall, the present study suggests that although the mutant binds inhibitors more effectively than the WT, with similar relative binding affinities, the post-binding structural and dynamical effects vary. Talazoparib inhibition has a notably stronger effect on the mutant compared to the WT. Specifically, under talazoparib inhibition, the mutant shows a more stable backbone of the catalytic domain (including the HD and ART domains), with notably stabilized N- and C-terminal residues, the H-Y-E triad, and the D-loop compared to the WT. However, higher destabilizing interactions are observed between the mutation site and HD domain under talazoparib inhibition than other inhibitors. This suggests that talazoparib provides the highest structural stability to PARP1’s catalytic domain while also enhancing destabilizing interactions between the HD domain and the mutation site among the inhibitors studied. Additionally, talazoparib inhibition significantly disrupts the functional fluctuations of terminal regions in the mutant, which are otherwise present in the WT. The primary motion of PARP1 remains unaffected by both inhibition and the mutation. Lastly, the limited understanding of the N-terminal’s role in PARP’s catalytic domain highlights opportunities for future research.

## Supporting information

Main draft and SI

## DATA AVAILABILITY

The input files for MD and TI-MD, .nmd files, movie of predominant motion of PARP1, and the trajectory for last 100 ns of a representative replica is provided in a Zenodo repository (DOI 10.5281/zenodo.14037435, https://doi.org/10.5281/zenodo.14037436).

## SUPPLEMENTARY DATA

Supplementary Data are available online.

### AUTHOR CONTRIBUTIONS

G.A.C. conceived the project, secured funding and resources. N.S. performed the MD simulations, and analyses. S.C. performed the TI simulations, EDA of non-bonded interactions with the mutation site, NMA, analyses and wrote the initial manuscript. G.A.C., N.S., and S.C wrote, edited, and approved the final version of the manuscript.

## ACKNOWLEDGEMENTS

This study was funded by NIH Grant No. R35GM151951. The computational time for this project was provided by the University of North Texas CASCaM CRUNTCh4 high-performance cluster, NSF Grant No. OAC-2117247, NSF ACCESS Project No. BIO240023, and the University of Texas at Dallas’ Cyberinfrastructure and Research Services, Ganymede and Titan HPC clusters.

## FUNDING

NIH R35GM191551; NSF OAC-2117247

## CONFLICT OF INTEREST

The authors declare no conflict of interests.

## Notes

https://doi.org/10.5281/zenodo.14037436

### Competing Interest Statement

The authors have declared no competing interest.

## REFERENCES

1. Harvey F. Lodish, A.B.C.K.M.K.M.P.S.A.B.H.L.P.P.T.M. (2008) Molecular cell biology sixth. W.H. Freeman, New York, ©2008.

2. Chatterjee, N. and Walker, G.C. (2017) Mechanisms of DNA damage, repair, and mutagenesis. Environ Mol Mutagen, 58, 235–263.

3. Rudolph, J., Jung, K. and Luger, K. (2022) Inhibitors of PARP: Number crunching and structure gazing. 10.1073/pnas.

4. Jeggo, P.A., Pearl, L.H. and Carr, A.M. (2016) DNA repair, genome stability and cancer: A historical perspective. Nat Rev Cancer, 16, 35–42.

5. Jackson, S.P. and Bartek, J. (2009) The DNA-damage response in human biology and disease. Nature, 461, 1071–1078.

6. Hoeijmakers, J.H.J. (2001) A plethora of damages in DNA The consequences of DNA injury Genome maintenance mechanisms for preventing cancer.

7. Yu, H., Ma, H., Yin, M. and Wei, Q. (2012) Association Between PARP-1 V762A Polymorphism and Cancer: Susceptibility: A meta-analysis. Genet Epidemiol, 36, 56–65.

8. Kim, M.Y., Zhang, T. and Kraus, W.L. (2005) Poly(ADP-ribosyl)ation by PARP-1: ‘PAR-laying’ NAD+ into a nuclear signal. Genes Dev, 19, 1951–1967.

9. Bulavko, E.S., Pak, M.A. and Ivankov, D.N. (2023) In Silico Simulations Reveal Molecular Mechanism of Uranyl Ion Toxicity towards DNA-Binding Domain of PARP-1 Protein. Biomolecules, 13.

10. Priyankha, S., Rajapandian, V., Palanisamy, K., Esther Rubavathy, S.M., Thilagavathi, R., Selvam, C. and Prakash, M. (2024) Identification of indole-based natural compounds as inhibitors of PARP-1 against triple-negative breast cancer: a computational study. J Biomol Struct Dyn, 42, 2667–2680.

11. Passeri, D., Camaioni, E., Liscio, P., Sabbatini, P., Ferri, M., Carotti, A., Giacchè, N., Pellicciari, R., Gioiello, A. and Macchiarulo, A. (2016) Concepts and Molecular Aspects in the Polypharmacology of PARP-1 Inhibitors. ChemMedChem, 10.1002/cmdc.201500391.

12. Rouleau, M., Patel, A., Hendzel, M.J., Kaufmann, S.H. and Poirier, G.G. (2010) PARP inhibition: PARP1 and beyond. Nat Rev Cancer, 10, 293–301.

13. Marchand, J.R., Carotti, A., Passeri, D., Filipponi, P., Liscio, P., Camaioni, E., Pellicciari, R., Gioiello, A. and Macchiarulo, A. (2014) Investigating the allosteric reverse signalling of PARP inhibitors with microsecond molecular dynamic simulations and fluorescence anisotropy. Biochim Biophys Acta Proteins Proteom, 1844, 1765–1772.

14. Lord, C.J. and Ashworth, A. PARP inhibitors: Synthetic lethality in the clinic.

15. Marchand, J.R., Carotti, A., Passeri, D., Filipponi, P., Liscio, P., Camaioni, E., Pellicciari, R., Gioiello, A. and Macchiarulo, A. (2014) Investigating the allosteric reverse signalling of PARP inhibitors with microsecond molecular dynamic simulations and fluorescence anisotropy. Biochim Biophys Acta Proteins Proteom, 1844, 1765–1772.

16. Kanev, P.B., Atemin, A., Stoynov, S. and Aleksandrov, R. (2024) PARP1 roles in DNA repair and DNA replication: The basi(c)s of PARP inhibitor efficacy and resistance. Semin Oncol, 51, 2–18.

17. Ménissier, J., Murcia, D.E., Niedergang, C., Trucco, C., Ricoul, M., Dutrillaux, B., Mark, M., Oliver, F.J., Masson, M., Dierich, A., et al. (1997) Requirement of poly(ADP-ribose) polymerase in recovery from DNA damage in mice and in cells (cellular response to DNA damage-raysalkylating agentsG2 arrestapoptosis).

18. Feltes, B.C. and Alvares, L. de O. (2024) PARP1 in the intersection of different DNA repair pathways, memory formation, and sleep pressure in neurons. J Neurochem, 10.1111/jnc.16131.

19. Huang, D. and Kraus, W.L. (2022) The expanding universe of PARP1-mediated molecular and therapeutic mechanisms. Mol Cell, 82, 2315–2334.

20. Spiegel, J.O., Van Houten, B. and Durrant, J.D. (2021) PARP1: Structural insights and pharmacological targets for inhibition. DNA Repair (Amst), 103.

21. Kanev, P.B., Atemin, A., Stoynov, S. and Aleksandrov, R. (2024) PARP1 roles in DNA repair and DNA replication: The basi(c)s of PARP inhibitor efficacy and resistance. Semin Oncol, 51, 2–18.

22. Pascal, J.M. (2018) The comings and goings of PARP-1 in response to DNA damage. DNA Repair (Amst), 71, 177–182.

23. Ménissier, J., Murcia, D.E., Niedergang, C., Trucco, C., Ricoul, M., Dutrillaux, B., Mark, M., Oliver, F.J., Masson, M., Dierich, A., et al. (1997) Requirement of poly(ADP-ribose) polymerase in recovery from DNA damage in mice and in cells (cellular response to DNA damage-raysalkylating agentsG2 arrestapoptosis).

24. Alemasova, E.E. and Lavrik, O.I. (2019) Poly(ADP-ribosyl)ation by PARP1: Reaction mechanism and regulatory proteins. Nucleic Acids Res, 47, 3811–3827.

25. Hurtado-Bagès, S., Knobloch, G., Ladurner, A.G. and Buschbeck, M. (2020) The taming of PARP1 and its impact on NAD+ metabolism. Mol Metab, 38.

26. Pushkarev, S. V., Kirilin, E.M., Švedas, V.K. and Nilov, D.K. (2024) Mechanism of PARP1 Elongation Reaction Revealed by Molecular Modeling. Biochemistry (Moscow), 89, 1202–1210.

27. Alemasova, E.E. and Lavrik, O.I. (2019) Poly(ADP-ribosyl)ation by PARP1: Reaction mechanism and regulatory proteins. Nucleic Acids Res, 47, 3811–3827.

28. Pushkarev, S. V., Kirilin, E.M., Švedas, V.K. and Nilov, D.K. (2024) Mechanism of PARP1 Elongation Reaction Revealed by Molecular Modeling. Biochemistry (Moscow), 89, 1202–1210.

29. Hurtado-Bagès, S., Knobloch, G., Ladurner, A.G. and Buschbeck, M. (2020) The taming of PARP1 and its impact on NAD+ metabolism. Mol Metab, 38.

30. Ali, A.A.E., Timinszky, G., Arribas-Bosacoma, R., Kozlowski, M., Hassa, P.O., Hassler, M., Ladurner, A.G., Pearl, L.H. and Oliver, A.W. (2012) The zinc-finger domains of PARP1 cooperate to recognize DNA strand breaks. Nat Struct Mol Biol, 19, 685–692.

31. Eustermann, S., Videler, H., Yang, J.C., Cole, P.T., Gruszka, D., Veprintsev, D. and Neuhaus, D. (2011) The DNA-binding domain of human PARP-1 interacts with DNA single-strand breaks as a monomer through its second zinc finger. J Mol Biol, 407, 149–170.

32. Kumar, C., Lakshmi, P.T.V. and Arunachalam, A. (2020) A mechanistic approach to understand the allosteric reverse signaling by selective and trapping poly(ADP-ribose) polymerase 1 (PARP-1) inhibitors. J Biomol Struct Dyn, 38, 2482–2492.

33. Bulavko, E.S., Pak, M.A. and Ivankov, D.N. (2023) In Silico Simulations Reveal Molecular Mechanism of Uranyl Ion Toxicity towards DNA-Binding Domain of PARP-1 Protein. Biomolecules, 13.

34. Marchand, J.R., Carotti, A., Passeri, D., Filipponi, P., Liscio, P., Camaioni, E., Pellicciari, R., Gioiello, A. and Macchiarulo, A. (2014) Investigating the allosteric reverse signalling of PARP inhibitors with microsecond molecular dynamic simulations and fluorescence anisotropy. Biochim Biophys Acta Proteins Proteom, 1844, 1765–1772.

35. van Beek, L., McClay, É., Patel, S., Schimpl, M., Spagnolo, L. and Maia de Oliveira, T. (2021) Parp power: A structural perspective on parp1, parp2, and parp3 in dna damage repair and nucleosome remodelling. Int J Mol Sci, 22.

36. Zandarashvili, L., Langelier, M.F., Velagapudi, U.K., Hancock, M.A., Steffen, J.D., Billur, R., Hannan, Z.M., Wicks, A.J., Krastev, D.B., Pettitt, S.J., et al. (2020) Structural basis for allosteric PARP-1 retention on DNA breaks. Science, eaax6367, 368.

37. Livraghi, L. and Garber, J.E. (2015) PARP inhibitors in the management of breast cancer: Current data and future prospects. BMC Med, 13.

38. Pommier, Y., O’connor, M.J. and De Bono, J. Laying a trap to kill cancer cells: PARP inhibitors and their mechanisms of action.

39. Jones, P., Wilcoxen, K., Rowley, M. and Toniatti, C. (2015) Niraparib: A Poly(ADP-ribose) polymerase (PARP) inhibitor for the treatment of tumors with defective homologous recombination. J Med Chem, 58, 3302–3314.

40. Wang, B., Chu, D., Feng, Y., Shen, Y., Aoyagi-Scharber, M. and Post, L.E. (2016) Discovery and Characterization of (8S,9R)-5-Fluoro-8-(4-fluorophenyl)-9-(1-methyl-1H-1,2,4-triazol-5-yl)-2,7,8,9-tetrahydro-3H-pyrido[4,3,2-de]phthalazin-3-one (BMN 673, Talazoparib), a Novel, Highly Potent, and Orally Efficacious Poly(ADP-ribose) Polymerase-1/2 Inhibitor, as an Anticancer Agent. J Med Chem, 59, 335–357.

41. Yu, J., Luo, L., Hu, T., Cui, Y., Sun, X., Gou, W., Hou, W., Li, Y. and Sun, T. (2022) Structure-based design, synthesis, and evaluation of inhibitors with high selectivity for PARP-1 over PARP-2. Eur J Med Chem, 227.

42. Rubiales-Martínez, A., Martínez, J., Mera-Jiménez, E., Pérez-Flores, J., Téllez-Isaías, G., Miranda Ruvalcaba, R., Hernández-Rodríguez, M., Mancilla Percino, T., Macías Pérez, M.E. and Nicolás-Vázquez, M.I. (2024) Design of Two New Sulfur Derivatives of Perezone: In Silico Study Simulation Targeting PARP-1 and In Vitro Study Validation Using Cancer Cell Lines. Int J Mol Sci, 25.

43. Bhatnagar, A., Nath, V., Kumar, N. and Kumar, V. (2024) Discovery of novel PARP-1 inhibitors using tandem in silico studies: integrated docking, e-pharmacophore, deep learning based de novo and molecular dynamics simulation approach. J Biomol Struct Dyn, 42, 3396–3409.

44. Mgoboza, C., Okunlola, F.O., Akawa, O.B., Aljoundi, A. and Soliman, M.E.S. (2022) Talazoparib Dual-targeting on Poly (ADP-ribose) Polymerase-1 and -16 Enzymes Offers a Promising Therapeutic Strategy in Small Cell Lung Cancer Therapy: Insight from Biophysical Computations. Cell Biochem Biophys, 80, 495–504.

45. Chadha, N. and Silakari, O. (2017) Identification of low micromolar dual inhibitors for aldose reductase (ALR2) and poly (ADP-ribose) polymerase (PARP-1) using structure based design approach. Bioorg Med Chem Lett, 27, 2324–2330.

46. Li, J., Zhou, N., Cai, P. and Bao, J. (2016) In silico screening identifies a novel potential PARP1 inhibitor targeting synthetic lethality in cancer treatment. Int J Mol Sci, 17.

47. Passeri, D., Camaioni, E., Liscio, P., Sabbatini, P., Ferri, M., Carotti, A., Giacchè, N., Pellicciari, R., Gioiello, A. and Macchiarulo, A. (2016) Concepts and Molecular Aspects in the Polypharmacology of PARP-1 Inhibitors. ChemMedChem, 10.1002/cmdc.201500391.

48. Priyankha, S., Rajapandian, V., Palanisamy, K., Esther Rubavathy, S.M., Thilagavathi, R., Selvam, C. and Prakash, M. (2024) Identification of indole-based natural compounds as inhibitors of PARP-1 against triple-negative breast cancer: a computational study. J Biomol Struct Dyn, 42, 2667–2680.

49. Lord, C.J. and Ashworth, A. PARP inhibitors: Synthetic lethality in the clinic.

50. Baptista, S.J., Silva, M.M.C., Moroni, E., Meli, M., Colombo, G., Dinis, T.C.P. and Salvador, J.A.R. (2017) Novel PARP-1 Inhibitor scaffolds disclosed by a dynamic structure-based pharmacophore approach. PLoS One, 12.

51. Turk, A.A. and Wisinski, K.B. (2018) PARP inhibitors in breast cancer: Bringing synthetic lethality to the bedside. Cancer, 124, 2498–2506.

52. Hu, H., Chen, B., Zheng, D. and Huang, G. (2020) Revealing the selective mechanisms of inhibitors to PARP-1 and PARP-2 via multiple computational methods. PeerJ, 2020.

53. Kamaletdinova, T., Fanaei-Kahrani, Z. and Wang, Z.Q. (2019) The enigmatic function of parp1: From parylation activity to par readers. Cells, 8.

54. Lord, C.J. and Ashworth, A. PARP inhibitors: Synthetic lethality in the clinic.

55. Kim, C., Chen, C. and Yu, Y. (2021) Avoid the trap: Targeting PARP1 beyond human malignancy. Cell Chem Biol, 28, 456–462.

56. Hopkins, T.A., Ainsworth, W.B., Ellis, P.A., Donawho, C.K., DiGiammarino, E.L., Panchal, S.C., Abraham, V.C., Algire, M.A., Shi, Y., Olson, A.M., et al. (2019) PARP1 trapping by PARP inhibitors drives cytotoxicity in both cancer cells and healthy bone marrow. Molecular Cancer Research, 17, 409–419.

57. Demin, A.A., Hirota, K., Tsuda, M., Adamowicz, M., Hailstone, R., Brazina, J., Gittens, W., Kalasova, I., Shao, Z., Zha, S., et al. (2021) XRCC1 prevents toxic PARP1 trapping during DNA base excision repair. Mol Cell, 81, 3018-3030.e5.

58. Hirota, K., Ooka, M., Shimizu, N., Yamada, K., Tsuda, M., Ibrahim, M.A., Yamada, S., Sasanuma, H., Masutani, M. and Takeda, S. (2022) XRCC1 counteracts poly(ADP ribose)polymerase (PARP) poisons, olaparib and talazoparib, and a clinical alkylating agent, temozolomide, by promoting the removal of trapped PARP1 from broken DNA. Genes to Cells, 27, 331–344.

59. Murai, J., Huang, S.Y.N., Das, B.B., Renaud, A., Zhang, Y., Doroshow, J.H., Ji, J., Takeda, S. and Pommier, Y. (2012) Trapping of PARP1 and PARP2 by clinical PARP inhibitors. Cancer Res, 72, 5588–5599.

60. Murai, J., Huang, S.Y.N., Das, B.B., Renaud, A., Zhang, Y., Doroshow, J.H., Ji, J., Takeda, S. and Pommier, Y. (2012) Trapping of PARP1 and PARP2 by clinical PARP inhibitors. Cancer Res, 72, 5588–5599.

61. Murai, J., Huang, S.Y.N., Renaud, A., Zhang, Y., Ji, J., Takeda, S., Morris, J., Teicher, B., Doroshow, J.H. and Pommier, Y. (2014) Stereospecific PARP trapping by BMN 673 and comparison with olaparib and rucaparib. Mol Cancer Ther, 13, 433–443.

62. Kamaletdinova, T., Fanaei-Kahrani, Z. and Wang, Z.Q. (2019) The enigmatic function of parp1: From parylation activity to par readers. Cells, 8.

63. Kumar, C., Lakshmi, P.T.V. and Arunachalam, A. (2020) A mechanistic approach to understand the allosteric reverse signaling by selective and trapping poly(ADP-ribose) polymerase 1 (PARP-1) inhibitors. J Biomol Struct Dyn, 38, 2482–2492.

64. Zhang, X., Wang, Y., Gari, A., Qu, C. and Chen, J. (2021) Pan-Cancer Analysis of PARP1 Alterations as Biomarkers in the Prediction of Immunotherapeutic Effects and the Association of Its Expression Levels and Immunotherapy Signatures. Front Immunol, 12.

65. Cashman, R., Zilberberg, A., Priel, A., Philip, H., Varvak, A., Jacob, A., Shoval, I. and Efroni, S. (2020) A single nucleotide variant of human PARP1 determines response to PARP inhibitors. NPJ Precis Oncol, 4.

66. Ravishankar, K., Jiang, X., Leddin, E.M., Morcos, F. and Cisneros, G.A. (2022) Computational compensatory mutation discovery approach: Predicting a PARP1 variant rescue mutation. Biophys J, 121, 3663–3673.

67. Murai, J., Huang, S.Y.N., Das, B.B., Renaud, A., Zhang, Y., Doroshow, J.H., Ji, J., Takeda, S. and Pommier, Y. (2012) Trapping of PARP1 and PARP2 by clinical PARP inhibitors. Cancer Res, 72, 5588–5599.

68. van Andel, L., Rosing, H., Zhang, Z., Hughes, L., Kansra, V., Sanghvi, M., Tibben, M.M., Gebretensae, A., Schellens, J.H.M. and Beijnen, J.H. (2018) Determination of the absolute oral bioavailability of niraparib by simultaneous administration of a 14C-microtracer and therapeutic dose in cancer patients. Cancer Chemother Pharmacol, 81, 39–46.

69. FDA approves maintenance treatment for recurrent epithelial ovarian, fallopian tube or primary peritoneal cancers (2017).

70. Jones, P., Wilcoxen, K., Rowley, M. and Toniatti, C. (2015) Niraparib: A Poly(ADP-ribose) polymerase (PARP) inhibitor for the treatment of tumors with defective homologous recombination. J Med Chem, 58, 3302–3314.

71. FDA approves rucaparib for maintenance treatment of recurrent ovarian, fallopian tube, or primary peritoneal cancer (2018).

72. Yubero, A., Estévez, P., Barquín, A., Sánchez, L., Santaballa, A., Pajares, B., Reche, P., Salvador, C., Manso, L., Márquez, R., et al. (2023) Rucaparib for PARP inhibitor-pretreated ovarian cancer: A GEICO retrospective subgroup analysis from the Spanish Rucaparib Access Program. Gynecol Oncol Rep, 48.

73. Yubero, A., Barquín, A., Estévez, P., Pajares, B., Sánchez, L., Reche, P., Alarcón, J., Calzas, J., Gaba, L., Fuentes, J., et al. (2022) Rucaparib in recurrent ovarian cancer: real-world experience from the rucaparib early access programme in Spain – A GEICO study. BMC Cancer, 22.

74. FDA approves talazoparib for gBRCAm HER2-negative locally advanced or metastatic breast cancer (2018) U.S. FOOD & DRUG.

75. Wang, B., Chu, D., Feng, Y., Shen, Y., Aoyagi-Scharber, M. and Post, L.E. (2016) Discovery and Characterization of (8S,9R)-5-Fluoro-8-(4-fluorophenyl)-9-(1-methyl-1H-1,2,4-triazol-5-yl)-2,7,8,9-tetrahydro-3H-pyrido[4,3,2-de]phthalazin-3-one (BMN 673, Talazoparib), a Novel, Highly Potent, and Orally Efficacious Poly(ADP-ribose) Polymerase-1/2 Inhibitor, as an Anticancer Agent. J Med Chem, 59, 335–357.

76. Papeo, G., Posteri, H., Borghi, D., Busel, A.A., Caprera, F., Casale, E., Ciomei, M., Cirla, A., Corti, E., D’Anello, M., et al. (2015) Discovery of 2-[1-(4,4-Difluorocyclohexyl)piperidin-4-yl]-6-fluoro-3-oxo-2,3-dihydro-1H-isoindole-4-carboxamide (NMS-P118): A Potent, Orally Available, and Highly Selective PARP-1 Inhibitor for Cancer Therapy. J Med Chem, 58, 6875–6898.

77. Thorsell, A.G., Ekblad, T., Karlberg, T., Löw, M., Pinto, A.F., Trésaugues, L., Moche, M., Cohen, M.S. and Schüler, H. (2017) Structural Basis for Potency and Promiscuity in Poly(ADP-ribose) Polymerase (PARP) and Tankyrase Inhibitors. J Med Chem, 60, 1262–1271.

78. Zandarashvili, L., Langelier, M.F., Velagapudi, U.K., Hancock, M.A., Steffen, J.D., Billur, R., Hannan, Z.M., Wicks, A.J., Krastev, D.B., Pettitt, S.J., et al. (2020) Structural basis for allosteric PARP-1 retention on DNA breaks. Science (1979), 368.

79. Ann-Gerd Thorsell, T.E.T.K.M.L.A.F.P.L.T.M.M.M.S.C. and H.S. (2017) Correction to “Structural Basis for Potency and Promiscuity inPoly(ADP-ribose) Polymerase (PARP) and Tankyrase Inhibitors”. J. Med. Chem., 60, 1262–1271.

80. Eric F Pettersen, T.D.G.C.C.H.G.S.C.D.M.G.E.C.M.T.E.F. (2004) UCSF Chimera--a visualization system for exploratory research and analysis. J Comput Chem ., 25, 1605–1612.

81. Pearlman, D.A., Case, D.A., Caldwell, J.W., Ross, W.S., Cheatham Iii, T.E., Debolt, S., Ferguson, D., Seibel, G. and Kollman, P. (1995) AMBER, a package of computer programs for applying molecular mechanics, normal mode analysis, molecular dynamics and free energy calculations to simulate the structural and energetic properties of molecules.

82. Case Ross C Walker, D.A. and Darden Junmei Wang Robert Duke, T.E. Amber 2018 Reference Manual Principal contributors to the current codes.

83. William L. Jorgensen; Jayaraman Chandrasekhar; Jeffry D. Madura; Roger W. Impey; Michael L. Klein (1983) Comparison of simple potential functions for simulating liquid water. J. Chem. Phys., 79, 926–935.

84. Maier, J.A., Martinez, C., Kasavajhala, K., Wickstrom, L., Hauser, K.E. and Simmerling, C. (2015) ff14SB: Improving the Accuracy of Protein Side Chain and Backbone Parameters from ff99SB. J Chem Theory Comput, 11, 3696–3713.

85. Gillespie, D.T. The chemical Langevin equation.

86. Nonlinear 8eneralized Langevin Equations.

87. Richard J. Loncharich, B.R.B.R.W.P. (1992) Langevin dynamics of peptides: The frictional dependence of isomerization rates of N-acetylalanyl-N’-methylamide. Biopolymers, 32, 523–535.

88. Götz, A.W., Williamson, M.J., Xu, D., Poole, D., Le Grand, S. and Walker, R.C. (2012) Routine microsecond molecular dynamics simulations with AMBER on GPUs. 1. generalized born. J Chem Theory Comput, 8, 1542–1555.

89. Kirkwood, J.G. (1935) Statistical mechanics of fluid mixtures. J Chem Phys, 3, 300–313.

90. Steinbrecher, T., Joung, I. and Case, D.A. (2011) Soft-core potentials in thermodynamic integration: Comparing one-and two-step transformations. J Comput Chem, 32, 3253–3263.

91. Roe, D.R. and Cheatham, T.E. (2013) PTRAJ and CPPTRAJ: Software for processing and analysis of molecular dynamics trajectory data. J Chem Theory Comput, 9, 3084–3095.

92. Humphrey, W., Dalke, A. and Schulten, K. (1996) VMD: Visual Molecular Dynamics.

93. Theses, M. and Stone, J.E. (1998) Scholars’ Mine Scholars’ Mine An efficient library for parallel ray tracing and animation An efficient library for parallel ray tracing and animation.

94. Sliarma, R. SpeechlGesture Interface to a.

95. Varshney Frederick Brooks, A.P. and William Wright, J. V (1994) Invited submission.

96. Eargle, J., Wright, D. and Luthey-Schulten, Z. (2006) Multiple Alignment of protein structures and sequences for VMD. Bioinformatics, 22, 504–506.

97. Frishman, D. and Argos, P. (1995) Knowledge-based protein secondary structure assignment. Proteins: Structure, Function, and Bioinformatics, 23, 566–579.

98. Sannerl, M.F., Olsonl, A.J. and Spehner2, J.-C. Fast and Robust Computation of Molecular Surfaces.

99. Stone, J.E., Gullingsrud, J. and Schulten, K. A System for Interactive Molecular Dynamics Simulation.

100. Dewage, S.W. and Cisneros, G.A. (2015) Computational analysis of ammonia transfer along two intramolecular tunnels in staphylococcus aureus glutamine-dependent amidotransferase (GatCAB). Journal of Physical Chemistry B, 119, 3669–3677.

101. Graham, S.E., Syeda, F. and Cisneros, G.A. (2012) Computational prediction of residues involved in fidelity checking for DNA synthesis in DNA polymerase i. Biochemistry, 51, 2569–2578.

102. Walker, A.R. and Cisneros, G.A. (2017) Computational Simulations of DNA Polymerases: Detailed Insights on Structure/Function/Mechanism from Native Proteins to Cancer Variants. Chem Res Toxicol, 30, 1922–1935.

103. Zandarashvili, L., Langelier, M.F., Velagapudi, U.K., Hancock, M.A., Steffen, J.D., Billur, R., Hannan, Z.M., Wicks, A.J., Krastev, D.B., Pettitt, S.J., et al. (2020) Structural basis for allosteric PARP-1 retention on DNA breaks. Science (1979), 368.

104. Langelier, M.-F., Planck, J.L., Roy, S. and Pascal, J.M. Structural Basis for DNA Damage-Dependent Poly(ADP-ribosyl)ation by Human PARP-1.

