## Supplementary material for "Impact of a Cancer-Associated Mutation on Poly(ADP-ribose) Polymerase1 Inhibition": Main draft and SI

Neel Shanmugam[1†], Shubham Chatterjee[2†], G. Andrés Cisneros[2,3,*]
[1] Department of Chemistry, University of North Texas, Denton, TX 76201, USA
[2] Department of Chemistry and Biochemistry, University of Texas at Dallas, Richardson, TX, 75080, USA
[3] Department of Physics, University of Texas at Dallas, Richardson, TX, 75080, USA

Present Address: Neel Shanmugam, Department of Computer Science, Columbia University, New York, New York, 10027, USA

[†]authors contributed equally to this work.

1. Root-Mean-Square Deviation (RMSD)

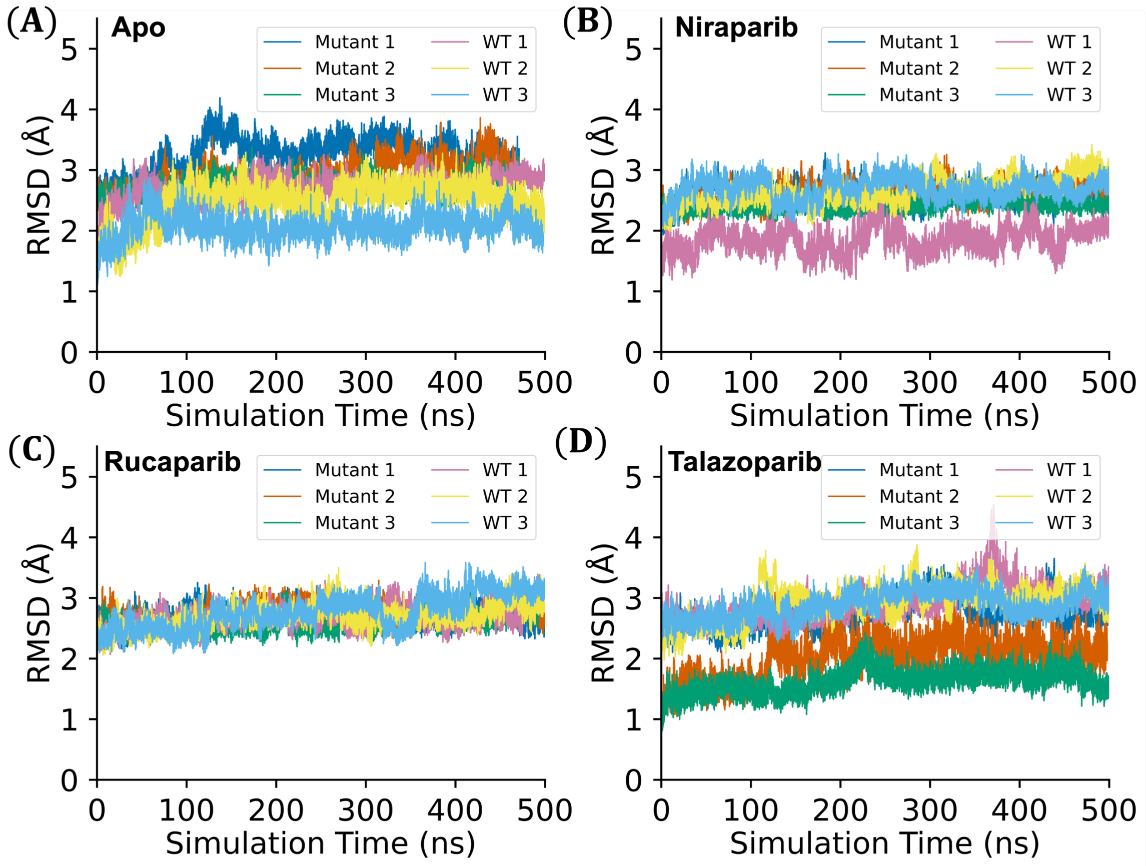


**Figure SI1.** Time dependent RMSD (in Å) of C$\alpha$ atoms across all the replicates (1,2, and 3) of WT and mutant systems under **(A)** apo **(B)** niraparib inhibition **(C)** rucaparib inhibition and **(D)** talazoparib inhibition

#### 2. Root-Mean-Square Fluctuation (RMSF)

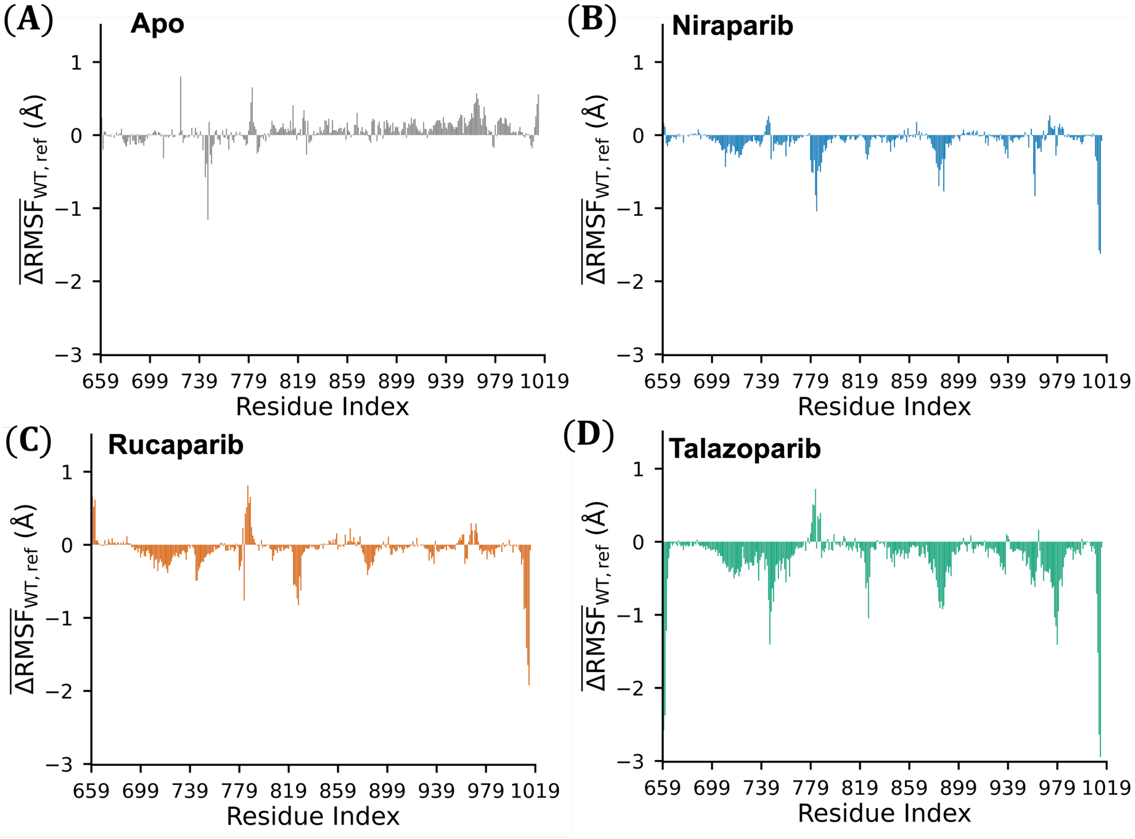


**Figure SI2.** Average change in RMSF (in Å) of PARP1 residues across all the replicates between WT and mutant with WT as reference under **(A)** apo **(B)** niraparib inhibition **(C)** rucaparib inhibition and **(D)** talazoparib inhibition

### 3. Energy Decomposition Analysis (EDA)

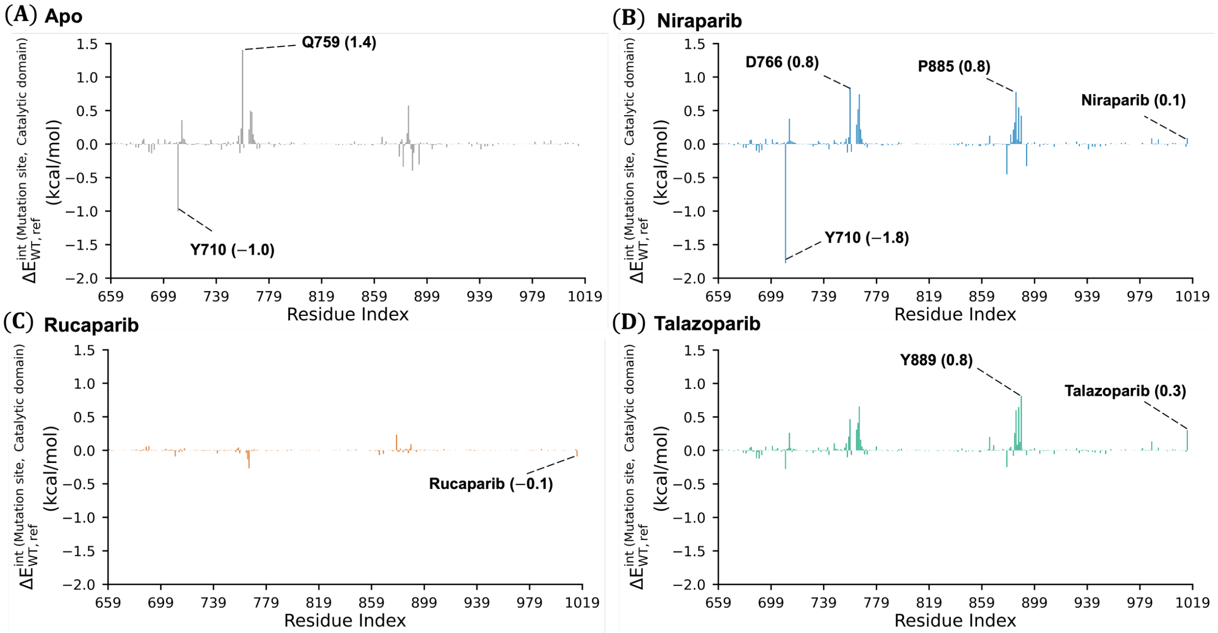


**Figure SI3.** Energy decomposition analysis of the nonbonded interactions of PARP1 residues and the respective inhibitor (expect for apo system) with the mutation site (V762 for WT and A762 for mutant) under **(A)** apo **(B)** niraparib-inhibition **(C)** rucaparib-inhibition and **(D)** talazoparib-inhibition. In all cases, the WT serves as the reference.

### 4. Thermodynamic Integration (TI)

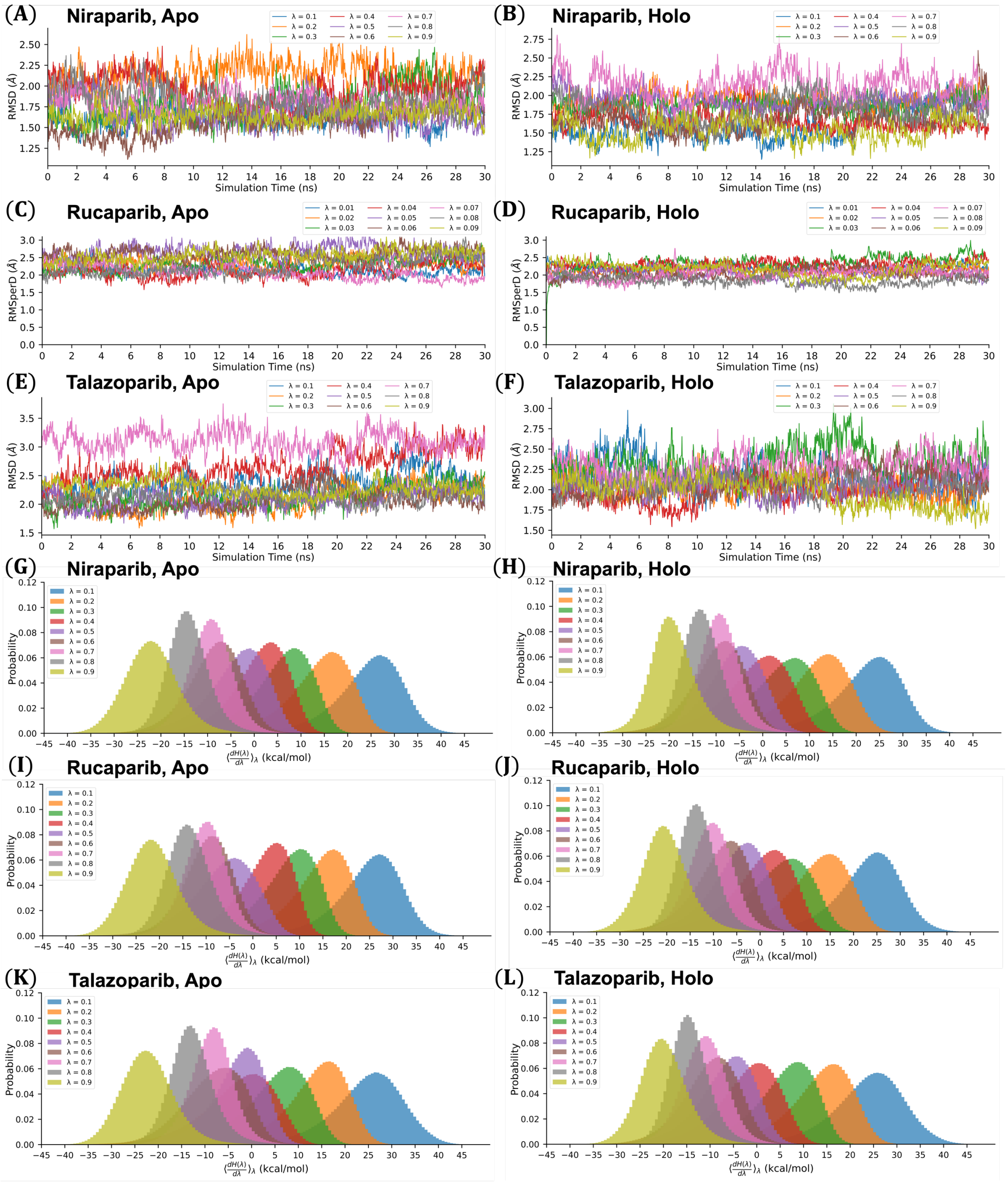


**Figure SI4.** Time dependent RMSD (in Å) of C$\alpha$ atoms during TI production for all λs for **(A)** apo structure (PDB ID: 4R6E) **(B)** holo structure with niraparib (PDB ID: 4R6E) **(C)** apo structure (PDB ID: 6VKK) **(D)** holo structure with rucaparib (PDB ID: 6VKK) **(E)** apo structure (PDB ID: 4UND) **(F)** holo structure with niraparib (PDB ID: 4UND). Probability distribution of λs for **(G)** apo structure (PDB ID: 4R6E) **(H)** holo structure with niraparib (PDB ID: 4R6E) **(I)** apo structure (PDB ID: 6VKK) **(J)** holo structure with rucaparib (PDB ID: 6VKK) **(K)** apo structure (PDB ID: 4UND) **(L)** holo structure with niraparib (PDB ID: 4UND).

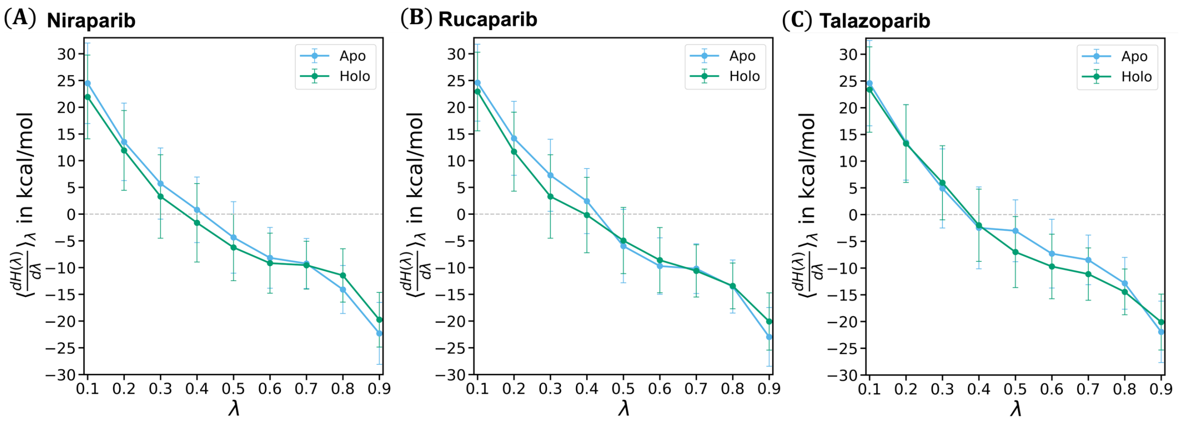


**Figure SI5.** $\langle \frac{\partial H(\lambda)}{\partial \lambda} \rangle_\lambda$ values (in kcal/mol) and corresponding standard deviation vs λ for apo and holo systems under **(A)** niraparib-inhibition **(B)** rucaparib-inhibition **(C)** talazoparib-inhibition. The difference in the areas under the curves (holo − apo) is the relative binding free energy of the inhibitor to the mutant compared to the WT.

### 5. Normal Mode Analysis (NMA)

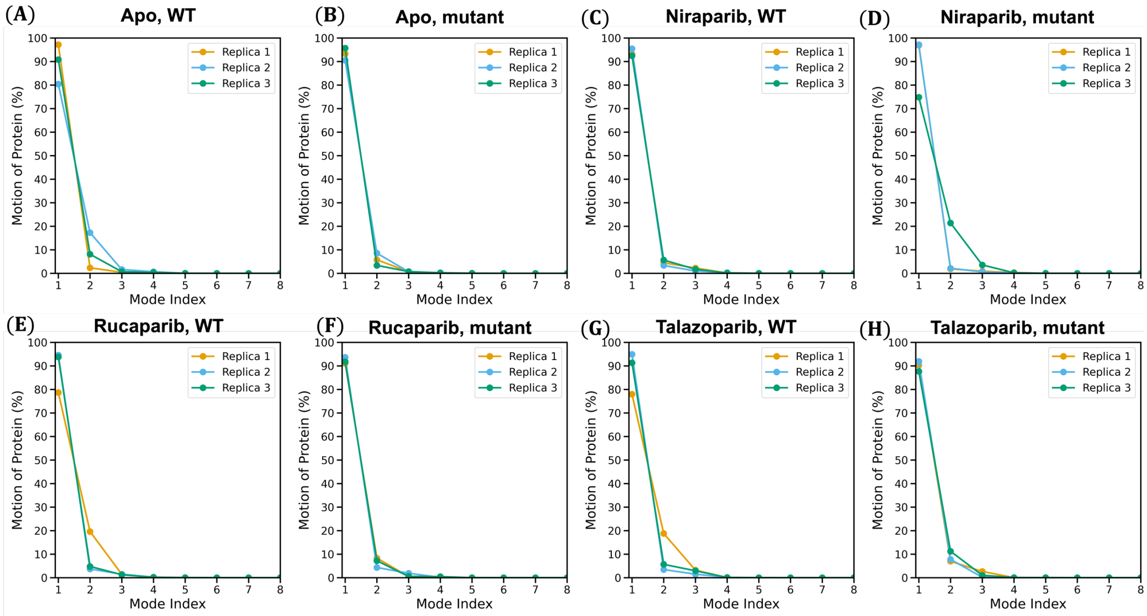


**Figure SI6. Normal Mode Analysis of WT and V762A mutant across three replicas**. Mode 1 of Normal Mode Analysis of (A) WT in apo system (B) mutant in apo system (C) WT under niraparib inhibition (D) mutant under niraparib inhibition (E) WT under rucaparib inhibition (F) mutant under rucaparib inhibition (G) WT under talazoparib inhibition (H) mutant under talazoparib inhibition.
